# Unsupervised Extraction of Epidemic Syndromes from Participatory Influenza Surveillance Self-reported Symptoms

**DOI:** 10.1101/314591

**Authors:** Kyriaki Kalimeri, Matteo Delfino, Ciro Cattuto, Daniela Perrotta, Vittoria Colizza, Caroline Guerrisi, Clement Turbelin, Jim Duggan, John Edmunds, Chinelo Obi, Richard Pebody, Ricardo Mexia, Ana Franco, Yamir Moreno, Sandro Meloni, Carl Koppeschaar, Charlotte Kjelsø, Daniela Paolotti

## Abstract

Seasonal influenza surveillance is usually carried out by sentinel general practitioners who compile weekly reports based on the number of influenza-like illness (ILI) clinical cases observed among visited patients. This practice for surveillance is generally affected by two main issues: i) reports are usually released with a lag of about one week or more, ii) the definition of a case of influenza-like illness based on patients symptoms varies from one surveillance system to the other, i.e. from one country to the other. The availability of novel data streams for disease surveillance can alleviate these issues; in this paper, we employed data from Influenzanet, a participatory web-based surveillance project which collects symptoms directly from the general population in real time. We developed an unsupervised probabilistic framework that combines time series analysis of symptoms counts and performs an algorithmic detection of groups of symptoms, hereafter called *syndromes*. Symptoms counts were collected through the participatory web-based surveillance platforms of a consortium called Influenzanet which is found to correlate with Influenza-like illness incidence as detected by sentinel doctors. Our aim is to suggest how web-based surveillance data can provide an epidemiological signal capable of detecting influenza-like illness’ temporal trends without relying on a specific case definition. We evaluated the performance of our framework by showing that the temporal trends of the detected syndromes closely follow the ILI incidence as reported by the traditional surveillance, and consist of combinations of symptoms that are compatible with the ILI definition. The proposed framework was able to predict quite accurately the ILI trend of the forthcoming influenza season based only on the available information of the previous years. Moreover, we assessed the generalisability of the approach by evaluating its potentials for the detection of gastrointestinal syndromes. We evaluated the approach against the traditional surveillance data and despite the limited amount of data, the gastrointestinal trend was successfully detected. The result is a real-time flexible surveillance and prediction tool that is not constrained by any disease case definition.

**Author summary:** This study suggests how web-based surveillance data can provide an epidemiological signal capable of detecting influenza-like illness’ temporal trends without relying on a specific case definition. The proposed framework was able to predict quite accurately the ILI trend of the forthcoming influenza season based only on the available information of the previous years. Moreover, we assessed the generalisability of the approach by evaluating its potentials for the detection of gastrointestinal syndromes. We evaluated the approach against the traditional surveillance data and despite the limited amount of data, the gastrointestinal trend was successfully detected. The result is a real-time flexible surveillance and prediction tool that is not constrained by any disease case definition.

## Introduction

Seasonal influenza is an acute contagious respiratory illness caused by viruses that can be easily transmitted from person to person. Influenza viruses circulate worldwide causing annual epidemics with the highest activity during winter seasons in temperate regions and produce an estimated annual attack rate of 3 to 5 million cases of severe illness and about 250 to 500 thousand deaths around the world [1]. National surveillance systems monitor the influenza activity through a network of general practitioners (GPs) who report the weekly number of influenza-like illness (ILI) cases among the overall patients [2]. These traditional surveillance systems for seasonal influenza are usually the primary source of information for healthcare officials and policymakers for monitoring influenza epidemics. However, classification of ILI cases in GPs reports is usually based on common clinical symptoms observed among patients and, as with any syndromic-based disease surveillance, case definitions of “influenza-like illness” can vary [3–7]. They typically include fever, cough, sore throat, headache, muscle aches, nasal congestion and weakness. There are also some works from hospital-based studies [8, 9], age-specific antiviral trials [4, 10, 11] and national surveillance activities [12] aimed at exploring suitable ILI case symptomatic descriptions but, so far, no unique definition has been widely adopted by the various national surveillance systems worldwide. For this reason, seasonal influenza surveillance in European countries remains rather fragmented. Only in recent years, some state members have adopted the case definition provided by the European Center for Disease Control and Prevention (ECDC) which defines an ILI case as the sudden onset of symptoms with one or more systemic symptoms (fever or feverishness, malaise, headache, myalgia) plus one or more respiratory symptoms (cough, sore throat, shortness of breath) [13]. Nevertheless, a significant fraction of European countries still adopts their own clinical case definition to compile seasonal influenza surveillance weekly reports.

In recent years the availability of novel data streams has given rise to a variety of non-traditional approaches for monitoring seasonal influenza epidemics [14–16]. Such new digital data sources can be exploited to capture additional surveillance signals that can be used to complement GPs surveillance data [17–20]. In this context, some so-called participatory surveillance systems have emerged in several countries around the world with the aim of monitoring influenza circulation through Internet reporting of self-selected participants [21–23]. One of these systems, the Influenzanet project [21], has been established in Europe since 2011 and it is now present in ten European countries. The system relies on the voluntary participation of the general population through a dedicated national website in each country involved in the project. Data are obtained on a weekly basis through an online survey [24] where participants are invited to report whether they experienced or not any of the following symptoms since their last survey: fever, chills, runny or blocked nose, sneezing, sore throat, cough, shortness of breath, headache, muscle/joint pain, chest pain, feeling tired or exhausted, loss of appetite, coloured sputum/phlegm, watery/bloodshot eyes, nausea, vomiting, diarrhoea, stomach ache, or other symptoms. Differently from most traditional surveillance systems, this participatory form of online surveillance allows the collection of symptoms in real time and directly from the general population, including those individuals who do not seek health care assistance. The list of proposed symptoms has been chosen to include the various ILI definitions adopted by national surveillance systems in Europe and, at the same time, to get a comprehensive list of symptoms that could be clearly articulated and understood by participants and would allow the detection of various circulating flu-related illnesses. Even though participatory systems generally suffer from self-selection bias, causing the sample to be non-representative of the general population [25], previous works have shown that web-based surveillance data can provide relevant information to estimate age-specific influenza attack rates [26, 27], influenza vaccine effectiveness [28], risk factors for ILI [29, 30], and to assess health care seeking behavior [31]. Moreover, it has been largely demonstrated that weekly ILI incidence rates that can be extracted from web-based surveillance data correlate well with ILI incidence as reported by GPs surveillance [27, 32, 33] (in such approaches, the number of ILI cases among the web platform participants was determined by applying the ECDC case definition [13] to the set of symptoms reported by participants).

An additional advantage of collecting symptoms directly from individuals among the general population in the various Influenzanet countries is that it is straightforward to compare the prevalence and the temporal dynamics of specific symptoms and/or groups of symptoms from one country to the other. In this work, we propose an approach which focuses on studying the temporal trends of groups of symptoms, hereafter called *syndromes*, as collected through the Influenzanet platforms. The goal is to develop a mathematical framework able to extract, in an unsupervised fashion, the groups of symptoms that are in good correlation with the ILI incidence as detected by traditional surveillance systems without imposing a priori a specific ILI case definition. By using the daily occurrence of symptoms (represented as boolean variables, with value 1 if a symptom is present and value 0 otherwise) as reported by the Influenzanet participants, it is possible to build a matrix whose rows are the weekdays during an influenza season and the columns are occurrences of symptoms as reported by the participants. Each matrix element thus corresponds to the number of times a specific symptom has been reported during a specific day of the influenza season. The result is a high-dimensional sparse data set from which meaningful features can be automatically extracted with the use of mathematical tools such as *Non-negative Matrix Factorization (NMF)* [34]. In particular, we are interested in extracting *latent*^1^ features of the matrix, corresponding to linear combinations of groups of symptoms, that are deemed as relevant by the NMF algorithm. By assuming that a specific combination of reported symptoms is the symptomatic expression of one or more illnesses experienced by the participants, i.e. of the *syndromes* affecting the individual, we can select those syndromes which better correlate with the traditional surveillance ILI incidence and adopt such linear combination of symptoms as the best approximation for the actual influenza-like illness circulating among the general population.

For this study, we employed data collected by Influenzanet platforms in several European countries over the course of six influenza seasons (from 2011-2012 to 2016-2017). We evaluated the performance of our method by comparing the emerged syndromes against the national surveillance data for the ILI incidence. The evaluation is performed both in a quantitative and in a qualitative way showing that the emerging syndromes closely follow the actual ILI incidence. The emergent syndrome consists of symptoms that are compatible with the traditional surveillance ILI case definition and stable for all the countries and can be employed to monitor fluctuations in the symptomatic expression of influenza across countries. We also assessed the predictive power of the proposed framework; in this case, excluding the data from the final season 2016-2017, we employed the models learnt on the previous seasons to decompose the symptoms counts for the season 2016-2017 predicting the forthcoming ILI syndrome. The predicted trend of the ILI component was quite accurate for all countries with correlations ranging from 0.60 to 0.85. As a final step, we evaluated the generalisation ability of this approach employing the proposed framework for the identification of gastrointestinal syndromes based on the same data stream of Influenzanet and comparing against the data from the traditional surveillance system. The overall encouraging results suggest that such methodology can be employed as a real-time flexible surveillance and prediction tool that is not constrained by any disease case definition which can be employed to monitor a wide range of symptomatic infectious diseases or to nowcast the influenza trend, devising public health communication campaigns.

## Materials and Methods

### Ethics Statement

This study was conducted in agreement with country-specific regulations on privacy and data collection and treatment. Informed consent was obtained from all participants enabling the collection, storage, and treatment of data, and their publication in anonymized, processed, and aggregated forms for scientific purposes. In addition, approvals by Ethical Review Boards or Committees were obtained, where needed according to country-specific regulations. In the United Kingdom, the Flusurvey study was approved by the London School of Hygiene and Tropical Medicine Ethics Committee (Application number 5530). In France, the Grippenet.fr study was approved by the Comité consultatif sur le traitement de l’information en matière de recherche (CCTIRS, Advisory committee on information processing for research, authorization 11.565) and by the Commission Nationale de l’Informatique et des Libertés (CNIL, French Data Protection Authority, authorization DR-2012-024). In Portugal, the Gripenet project was approved by the National Data Protection Committee and also by the Ethics Committee of the Instituto Gulbenkian de Ciência.

### Data Collection

Since the winter season of 2011-2012, the Influenzanet platforms share a common and standardized data collection approach throughout the nine European countries involved, namely: Belgium (BE), Denmark (DK), France (FR), Ireland (IE), Italy (IT), Netherlands (NL), Portugal (PT), Spain (ES) and United Kingdom (UK). In each of the Influenzanet countries, the national platform is coordinated by a team of local researchers from Universities, Research Institutions or Public Health Institutions and consists of a website where individuals can register and have access to a personal account where they can insert and update their data. The platforms are disseminated among the general population through press releases, general media campaigns, specific dissemination events (e.g. science fairs) or word of mouth. Participation is voluntary and anonymous and all the residents of the participating countries can enrol. Upon registration, individuals are asked to complete an online Intake Questionnaire covering basic questions such as age, gender, household size and composition, home location, workplace, etc. [35]. Participants can also create accounts on behalf of other members of their family or household, thus enabling, for instance, parents to record data for their children. Registered participants are then reminded weekly, via an e-mail newsletter, to fill in a Symptoms Questionnaire [35] in which they are presented with a list of general, respiratory and gastrointestinal symptoms (18 in total, reported in Table 1) and asked whether since the last time they visited the platform they experienced any symptoms among those listed. In this study, we analyzed the data collected by the various national platforms from November 2011 to April 2017.

Seasonal influenza is traditionally monitored by national networks of general practitioners (GPs) who report the weekly number of visited patients with influenza-like illness symptoms according to the national ILI case definition. Despite some practical limitations, mainly due to an heterogeneous population coverage and to considerable delay in disseminating data, such traditional surveillance data are generally considered as ground truth. In this study, we collected the weekly ILI incidence data for 6 influenza seasons, from 2011-2012 to 2016-2017, from the ECDC dedicated web page [36] for all countries, expect France, for which, instead, we obtained the weekly data on the ILI incidence and gastrointestinal infections directly from the national network, called *Réseau Sentinelles* [37]. All reports were accessed and downloaded in March 2017.

**Table 1.** Symptoms included in the weekly questionnaire. List of the 18 symptoms presented to participants in the weekly Symptoms Questionnaire, plus the sudden onset, i.e. if symptoms appeared suddenly over a few hours.

|  |  |  |  |
| --- | --- | --- | --- |
| Fever | Chills | Runny/blocked nose | Sneezing |
| Sore throat | Cough | Shortness of breath | Headache |
| Muscle/joint pain | Chest pain | Feeling tired (malaise) | Loss of appetite |
| Coloured Sputum/<br>Phlegm | Watery, bloodshot eyes | Nausea | Vomiting |
| Diarrhoea | Stomachache | Sudden Onset |  |

### Data Preprocessing

In general, inclusion criteria of participants in the data analysis vary depending on the specific aim of the study [25, 33, 38, 39]. In our case, we included only those individuals who are registered on the national platforms and have filled in at least one survey per season. Specifically, we consider only one survey per participant for each week, keeping the last one if multiple surveys were submitted during the same week. This choice corresponds to a loss of approximately 5% of the total number of surveys and will allow assuming that symptoms reported in a week are proportional to the number of participants in that week. Moreover, to reduce the noise due to low participation rates at the beginning of the influenza season, we included in the analysis only those weeks in which the number of surveys corresponded at least to 5% of the total number of the surveys filled during the week with the highest participation for that season. In S1 Table in the supporting information, we present some descriptive statistics for each country, including: (i) the number of seasons available, (ii) the average number of participants per season, (iii) the average number of weekly surveys per season, (iv) the average percentage of surveys with at least one symptom, and (v) the average number of surveys per participant per season.

### Temporal Syndrome Modeling and Non-negative Matrix Factorization

In this section, we describe the methodology employed to extract the latent features from the self-reported symptoms collected by the various Influenzanet platforms of the participating countries. Our approach relies on the assumption that a specific group of self-reported symptoms corresponds to the symptomatic expression of one or more illnesses, hereafter called *syndromes*, circulating among the population sample participating in the study. In our study we consider a total of 19 symptoms, corresponding to the 18 symptoms presented in the weekly Symptoms Questionnaire plus an additional symptom, called “Sudden onset”, referring to the sudden appearance of symptoms, typically over the course of the previous 24 hours (see Table 1). Symptoms are treated as binary boolean variables having value 1 if the symptom is present and 0 if the symptom is absent.; then, we aggregated across all participants to build a matrix **X** = [*x_ij_*], whose elements contain the occurrences of each symptom *j* ∈ {1*, .., J*} during each day *i* ∈ {1*, .., I*}. In other words, each element of the matrix corresponds to the number of times each symptom has been reported on each day of the influenza seasons under study. The result is a high-dimensional sparse matrix that can be linearly decomposed through a Non-negative Matrix Factorizations (NMF) technique [34]. We opted for NMF since its non-negativity constraint offers the advantage of a straightforward interpretation of the results as positive quantities that can then be associated with the initial symptoms. This approach is equivalent to a “blind source separation” problem [40] in which neither the sources nor the mixing procedure is known, but only the resulting mixed signals are measured. In our case, the time series corresponding to the daily symptoms counts are measured by the Influenzanet platforms and can be considered as the result of a linear mixing process driven by unknown sources, i.e. the latent syndromes. In the following we will use interchangeably the terms *syndrome*, *source* or *component*. According to this consideration, each element *x_ij_* of the matrix **X** can be expressed as follows:

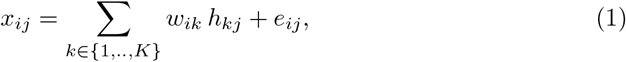

where the coefficients *h_kj_* describe the set of the unknown *K* sources, the factor *w_ik_* represents the time-dependent mixing coefficients, and the terms *e_ij_* correspond to the approximation error. The mixing equations 1 can be equivalently expressed in matrix notation as:

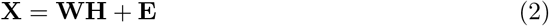

where:

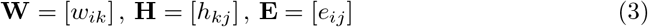

It is worth stressing that in this representation the matrix **X** is known, while the matrices **W** and **H** are unknown and determined by the NMF algorithm. In particular, we used a variation of the NMF algorithm that minimizes the Kullback-Leibler divergence loss function [41] defined as follows:

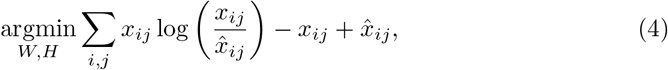

where:

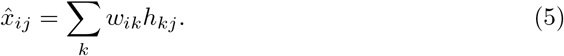

To minimize this function, we adopted the multiplicative update rules described in [42]. Note that different initializations of the matrices **W** and **H** might lead to different local minima, making the interpretation of the results not straightforward. To overcome this issue, we use an initialization technique called Non-negative Double Singular Value Decomposition [43], that is based on a probabilistic approach equivalent to the probabilistic latent semantic analysis (pLSA) [44], employed in the context of semantic analysis of text corpora. Since the two approaches of NMF and pLSA are equivalent (see [45] for more details), the results of our matrix decomposition can be probabilistically interpreted as a mixture of conditionally independent multinomials, that we call *p*(*i, j*). We can then write:

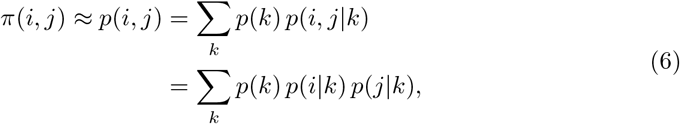

where:

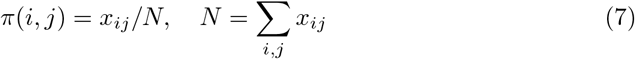

and *N* is the total number of symptoms counts.

According to 6, the total number of symptoms counts will be proportionally split among *K* latent sources according to *p*(*k*), which is the probability to observe a specific component *k*; *p*(*i*|*k*) is the probability to observe a component *k* in a day *i* and *p*(*j*|*k*) is the probability to observe a specific symptom *j* in a component *k*, and they can be expressed as follows:

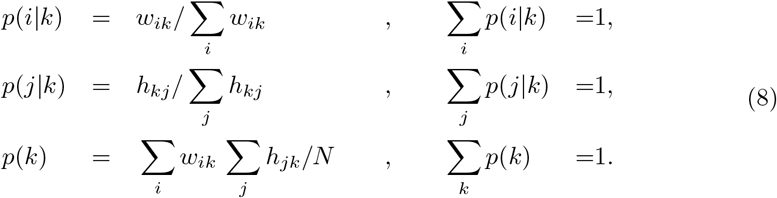

At this point, 8 allows to determine the probability *p*(*i, k*) that, rescaled on the total number of symptoms counts *N*, yields the desired decomposition procedure, *y_ik_*, which represents the contribution of a specific component *k* in a day *i*, given by the following expression:

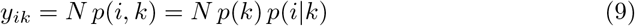

Thus, the final step in our approach is to determine the optimal number of components *k_min_* to be used for the decomposition. A natural upper bound for *k* would be the total number of symptoms, i.e. 19. We need to determine the number of components with the best tradeoff between a model that best approximates the original matrix *X* and at the same time does not overfit the data. Each time we minimize the loss function 4 for a specific number of components *k*, we obtain a candidate decomposition.

To determine the best decomposition, we use an approximated model selection criterion, known as the Akaike Information Criterion [46]. In particular, we employ the corrected version of the Akaike Information Criterion (*AIC*) proposed in [47], valid for finite sample sizes. For each of the candidate decompositions generated by the various values of *k*, we estimate the AIC criterion, *AIC_k_*, expressed as:

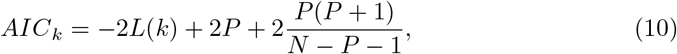

where, *L*(*k*) is the log-likelihood of the model with *k* components, defined in [45] as:

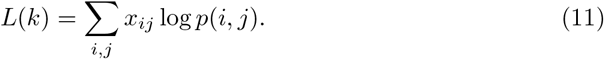

*P*, is the number of parameters of the model with:

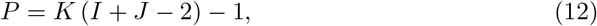

where *K* is the upper bound for the number of components, *I* is the total number of days and *J* is the total number of symptoms. The best candidate decomposition is the one that minimizes 10 and we denote as *AIC_k_min__*. The final result is a model, that we call *yik_min_* consisting of *k_min_* components that best apporoximate the original matrix *X*.

## Data Analysis

We applied the aforementioned framework to the data collected by the Influenzanet platforms in nine European countries over the course of six influenza seasons (from 2011-2012 to 2016-2017). For each country, we applied the decomposition algorithm to the symptoms’ matrix **X** as represented in 2 and, based on the AIC criterion, we obtain the “optimum” number of components, *k_min_*, for the decomposition.

Among the *k_min_* latent components, or syndromes, extracted for each country, we identify by means of Pearson correlation the one that shows a trend similar to the time-series recorded by the sentinel-based surveillance for influenza-like illness. In the following, we denote this component as IN_NMF. This component will correspond to the combination of symptoms that more closely represent the ILI time series recorded by the traditional surveillance, and hence, it can be used to build a data-driven, unsupervised ILI case definition, which is the ultimate goal of this study. We performed a weekly aggregation of the daily counts of the emerged syndromes so that we can compare them directly with the weekly incidence reported by the traditional surveillance of sentinel doctors.

To estimate the quality of the IN_NMF signal selected for each country, we also assessed the Pearson correlation between: (i) IN_NMF and the time series obtained by applying the ECDC ILI definition on the Influenzanet data (hereafter called IN ECDC); (ii) the IN NMF and the ILI incidence as reported by the national surveillance systems per country (hereafter called GP); and (iii) the IN ECDC and the ILI incidence reported by the national surveillance systems per country (GP). The reported correlations refer to the time series over the entire period (2011-2017).

As a final step, we explored the predictive power of the proposed methodology by applying the decomposition framework only on data collected from 2011 to 2016 (training phase) and predicting the ILI incidence signal for the season 2016-2017 (testing phase). To test the validity of this predicted signal, we have evaluated the Pearson correlation of the predicted time series for 2016-2017 and the ILI incidence reported by the national surveillance systems per country (GP). Moreover, we calculated the Pearson correlation between ILI incidence obtained by applying the ECDC case definition to raw Influenzanet data (IN ECDC) and the respective predicted time series, IN NMF, for the season 2016-2017.

To assess the generalisation potential of our framework in identifying syndromes not related to ILI (e.g. respiratory versus gastrointestinal), we used it to identify the syndrome related to gastrointestinal episodes by performing the Pearson correlation with data provided by the traditional official surveillance in France^2^. The identified component is denoted as IN_Gastro.

## Results and Discussion

### Selection of Components

Fig. S1 in the supporting information, depicts an exploration on the relative AIC values of a series of candidate models (*AIC_c_*(*k*) *− AIC_c_*(*k_min_*), with *k* ∈ [1, 6]), estimated according to 10. For the majority of the countries, the optimal decomposition consisted of *k_min_* = 2 components, with the exceptions of the Netherlands with *k_min_* = 3, Belgium with *k_min_* = 3 and France with *k_min_* = 4. Figures S2, S3, S4 and S5 in the supporting information depict the respective time series of all the emerging *k_min_* components and the symptomatic composition for each country. The component which, according to our framework, expresses the ILI incidence (IN_NMF) for each country is highlighted by a blue square.

### ILI Component Analysis

In the left panel of Fig. 1, the IN_NMF component for each country is shown in comparison to the ILI signal as recorded by the traditional surveillance, GP. To allow for visual comparison, the time series of the IN_NMF component was rescaled on the traditional surveillance GP time series with a fixed scaling factor. In the right panel of Fig. 1, the break-down of symptoms for each country’s IN_NMF component is expressed in terms of probabilistic contributions, denoted as *p*(*j*|*k*), as described in 6. In terms of symptoms’ composition, IN_NMF appears to be stable across the various countries and consistent with the expected set of symptoms clinically associated with ILI. The top contributing symptoms are fever, chills and feeling tired, often reported in combination with a sudden onset of symptoms. Notably, all these symptoms contribute for more than 10% to the overall component composition. This is consistent across all the countries and it is the most important result of this study since it represents the basis towards the development of a common ILI definition. Small heterogeneities in the component composition across countries are most likely due to differences in the ILI case definitions used by sentinel doctors in each country which are reflected in the data that we use as ground-truth. In principle, this issue could be overcome in the future by using as ground truth seroprevalence data instead than ILI incidence as captured by sentinel doctors which use different ILI definitions.

**Fig 1.**
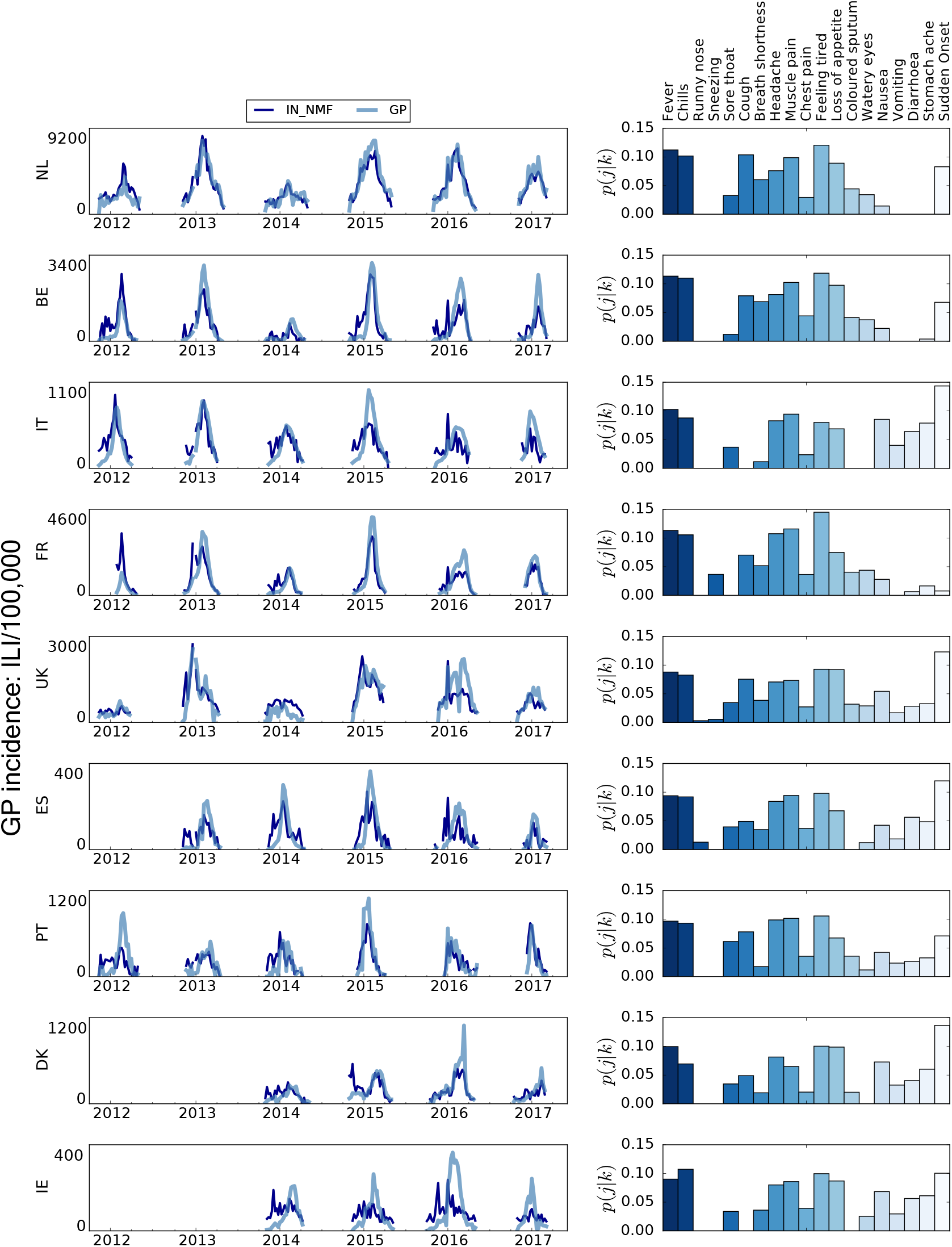
Left panel: qualitative comparison between the IN NMF and the national surveillance incidence (GP) time series. To allow for an easier visual inspection the depicted IN NMF syndromes are rescaled by a fixed factor to the respective GP incidence. On the y-axis, the sample size of the GP incidence is reported. Right panel: contribution of each symptom to the automatically selected IN_NMF component. The bars are coloured for readability purposes only.

### Model evaluation

Table 2 reports all the correlations mentioned in the Data Analysis section. For all countries, the correlation between the IN NMF component and the IN_ECDC is very high, more than 0.82 for all cases, showing that the IN NMF signal captures symptoms highly compatible with those present in the ECDC ILI definition applied to the Influenzanet data. However, carefully examining rows (ii) and (iii) of Table 2, we note slight variations per country. For the Netherlands, Belgium, and Ireland the IN NMF components are a much better fit (higher correlations) to the incidence data (GP) with respect to the time series obtained by applying the traditional ILI definition on the Influenzanet data (IN_ECDC). For the UK, Spain, Denmark, and Portugal the IN NMF components perform equally well as the IN_ECDC. For Italy and France, the IN NMF component had a slightly lower correlation (about 11% and 7% less respectively) with the ECDC surveillance data than the IN_ECDC. Ireland is the only country for which we obtain a low correlation between the surveillance and IN_ECDC probably due to the limited number of participants. In spite of this, we note that IN NMF performs much better than the IN_ECDC in capturing the incidence trend. This variation in performance is not an issue for the goal of this work since our focus is on paving the way towards a common cross-country ILI definition rather than finding the perfect signal that correlates best with the national traditional surveillance and the loss in performance of IN_NMF with respect to IN_ECDC for Italy and France is only a small percentage. *Prediction Evaluation:* The results of the prediction analysis described in the Data Analysis section are shown in the fourth row of Table 2 (iv), which reports the correlations of the predicted IN_NMF syndrome and the national surveillance for the season 2016-2017 (GP). The correlation between the two time series is good, ranging from 0.60 to 0.85, for all the countries. We also report, in Table 2 (v), the correlation between the IN_NMF predicted component and the time series emerged from applying the ECDC definition on the Influenzanet data for the season (2016-2017). Also in this case, the predicted trend of the ILI component had high correlations, ranging from 0.60 to 0.85 ( Table 2 (iv)).

**Table 2.**
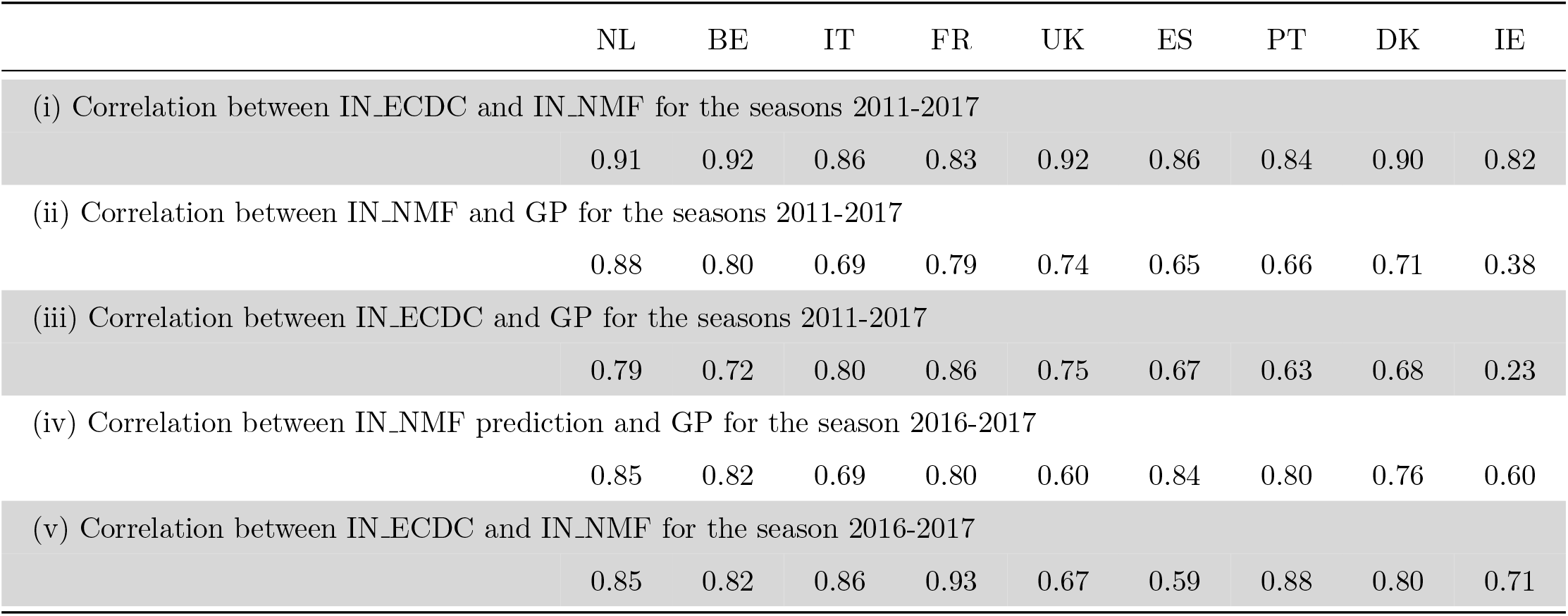
(i) Pearson correlation between the time series of IN_NMF with the respective time series produced when applying the ILI definition on the Influenzanet data (IN_ECDC). The two signals are highly correlated for all countries. (ii) Pearson correlation between IN NMF and the respective ILI incidence reported by the national surveillance systems per country (GP). iii) Pearson correlation between ILI incidence obtained by applying the ECDC case definition to raw Influenzanet data IN_ECDC) and ILI incidence reported by the national surveillance systems per country (GP). (iv) Pearson correlation between the predicted 2016-2017 IN_NMF and ILI incidence reported by the national surveillance systems per country (GP) for the season 2016-2017. (v) Pearson correlation between ILI incidence obtained by applying the ECDC case definition to raw Influenzanet data (IN_ECDC) and the respective IN_NMF for the 2016-2017. Note that the reported correlations are not averages per ILI seasons per country but the correlation of the time series of the entire period (2012-2017 for (i),(ii) and (iii) and 2016-2017 for (iv) and (v)) between the IN NMF and the respective GP time series for each country.

### Gastro Component Evaluation

In the left panel of Fig. 2, we show the time series for the incidence of acute diarrhoea episodes (*GP Gastro*) as detected by the official national surveillance in France, and the time series of the syndrome identified by our framework (*IN Gastro*). The Pearson correlation between the extracted syndrome and the official surveillance data is *ρ* = 0.66. In the right panel of Fig. 2 we depict the probabilistic contribution of each symptom to the IN Gastro syndrome. Emerging symptoms, in this case, include also stomach ache, diarrhoea, and vomiting, which are in line with our expectations. Even if respiratory symptoms like runny nose or sneezing are present, the contribution of fever and chills (which were the main contributors to the IN_NMF signal) is almost negligible. This corresponds to a rather good capability of the framework in discriminating between diverse syndromes. Despite limitations of the data availability, these preliminary findings indicate that the remaining latent components of the decomposition may express syndromes related to allergies, common-cold or gastroenteritis. Understandably, adequate surveillance data are required to make a firm statement and reach a robust interpretation of the syndromes.

**Fig 2.**
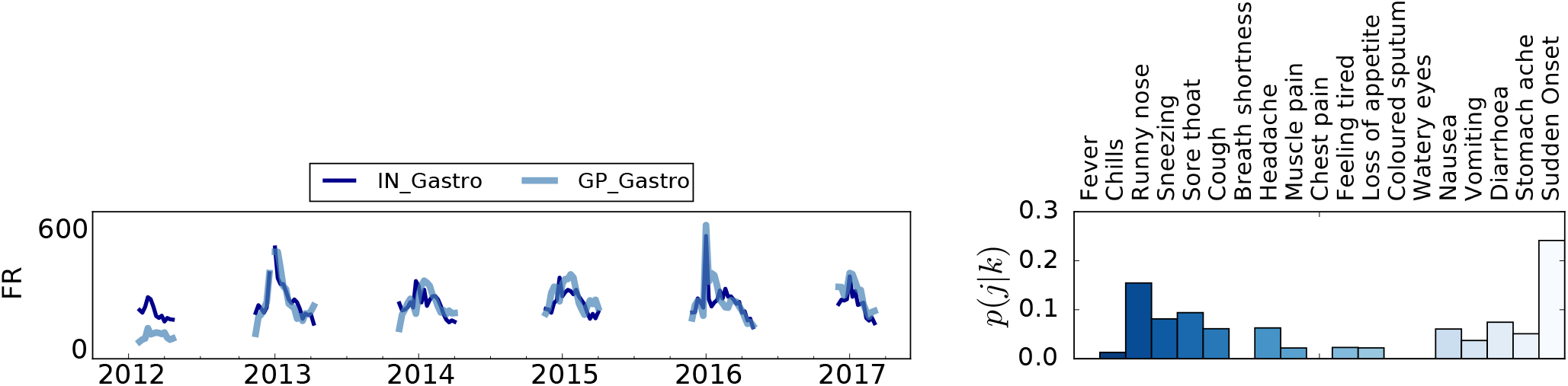
Left panel: Time series comparison between IN_Gastro component and the national surveillance data (GP_Gastro) for France. To allow for an easier visual inspection the depicted IN_Gastro syndrome is rescaled by a fixed factor on the respective GP_Gastro incidence. On the y-axis, the sample size of the GP incidence is reported. Right panel: symptomatic contribution of the automatically selected IN_Gastro component. The bars are colored for readability purposes only.

### Limitations

The limitations of this approach are two-fold; first, it is based on syndromic data and thus the specificity with respect to measuring the actual circulation of influenza viruses among the population is relatively low. We could overcome this issue with the integration of data from virologically confirmed cases in the analytical framework. Second, information provided by volunteers about self-reported symptoms might be affected by cognitive biases in assessing and recalling the symptoms. This is an unavoidable potential issue in the symptoms reporting which we cannot account for in our framework.

## Conclusions

The practice of seasonal influenza surveillance is generally affected by the fact that case definitions for influenza-like illness based on clinical symptoms are heterogeneous and might vary across different surveillance systems and hence across different countries. To overcome this issue, we propose an unsupervised probabilistic framework based on self-reported symptoms collected daily through a network of participatory Web-based influenza surveillance platforms in Europe. The approach, which relies on a Non-negative matrix factorization of the daily symptoms matrix, is capable to produce an epidemiological signal that does not rely on a specific a priori case definition and that follows closely the temporal trend of influenza-like illness as detected by sentinel doctors surveillance. The emerging signal, compared against national surveillance data, successfully captures the ILI incidence trend for all countries included in this study. Moreover, the probabilistic contributions of the symptoms in the overall composition of the emerging signal are stable across countries, paving the way towards the development of a common data-driven ILI case definition. We also demonstrate that the proposed approach can be employed to predict the forthcoming ILI incidence and we tackle the generalisation abilities of this approach by using it to identify gastrointestinal syndromes. We can thus conclude that there is great potential in using symptoms directly collected from the general population to inform unsupervised algorithmic approaches aimed at detecting circulating bouts of illnesses without imposing an a priori case definition. The standardized technological and epidemiological framework and the ability to monitor symptoms from the general population, including individuals who don’t seek medical assistance, provided by the Influenzanet participatory surveillance platforms are what enable the application of unsupervised algorithmic approaches such as the one presented in this work. In the next future, we will include data from virologically confirmed influenza cases as ground truth to enhance the specificity of our framework. Moreover, the flexibility provided by the participatory surveillance platforms in terms of symptoms that can be collected from the general population enable the possibility to extend the framework to other diseases, provided that traditional surveillance data are available to train the framework.

## Acknowledgments

The authors declare no competing financial interests. D.Pa and D.Pe. acknowledge support from H2020 FETPROACT-GSS CIMPLEX Grant No. 641191. K.K, M. D., C.C, D.Pa., and D.Pe. acknowledge support from the “Lagrange Project” of the ISI Foundation funded by the Fondazione CRT. Y.M. acknowledges support from the Government of Araǵon, Spain through a grant to the group FENOL and by MINECO and FEDER funds (grant FIS2017-87519-P). The authors would like to thank all the participants who took part in the participatory surveillance systems over the past years.

## Author Contributions

MD,KK,DPa,CC conceived the idea. MD,KK analyzed the data. DPa,VC,JD,JE,RM,YM,CK,CK provided the data for each participating country. KK,MD,DPa,DPe wrote the paper. All authors reviewed the paper.

## Supporting Information

**S1 Table.** Descriptive statistics regarding the available Influenzanet data for each country.

|  | NL | BE | IT | FR | UK | ES | PT | DK | IE |
| --- | --- | --- | --- | --- | --- | --- | --- | --- | --- |
| Number of seasons | 6 | 6 | 6 | 6 | 6 | 5 | 6 | 4 | 4 |
| Average participants per season | 13,450 | 4,209 | 1,830 | 5,757 | 4,676 | 526 | 1,663 | 1,391 | 406 |
| Average # surveys per season | 206,987 | 67,420 | 17,807 | 68,567 | 45,543 | 5,894 | 17,852 | 22,782 | 3,220 |
| Average % of surveys with symptoms | 20% | 16% | 19% | 20% | 29% | 22% | 17% | 18% | 25% |
| Average of surveys per participant per season | 15 | 16 | 9 | 12 | 9 | 11 | 10 | 16 | 8 |

### S2 Appendix. AIC Criterion

For each country, we generated a series of models by increasing the number of syndromes with *k* ∈ [1, 6]. For each of the models, we estimated the AIC criterion, *AIC_c_*(*k*) (Eq. 10). The best model was the one to minimize the Eq. 10, denoted as *AIC_c_*(*k_min_*), consisting of *k_min_* syndromes. Fig. S1 depicts, for each country, the relative likelihood (*AIC_c_*(*k*) − *AIC_c_*(*k_min_*)) of each candidate model with *k* syndromes (*AIC_c_*(*k*)), as compared to the model with the minimum *AIC_c_* score (*AIC_c_*(*k_min_*)).

### S3 Appendix Complete Set of the Extracted Latent Components

Comparative Analysis of the consistency and time series of the amount *y_ik_* which refers to the total number of counts associated to a syndrome *k* in the day *i* for all the emerged syndromes for each country. The green box indicates selected IN_NMF by the algorithm.

**Supplemental Information, Figure S1.**
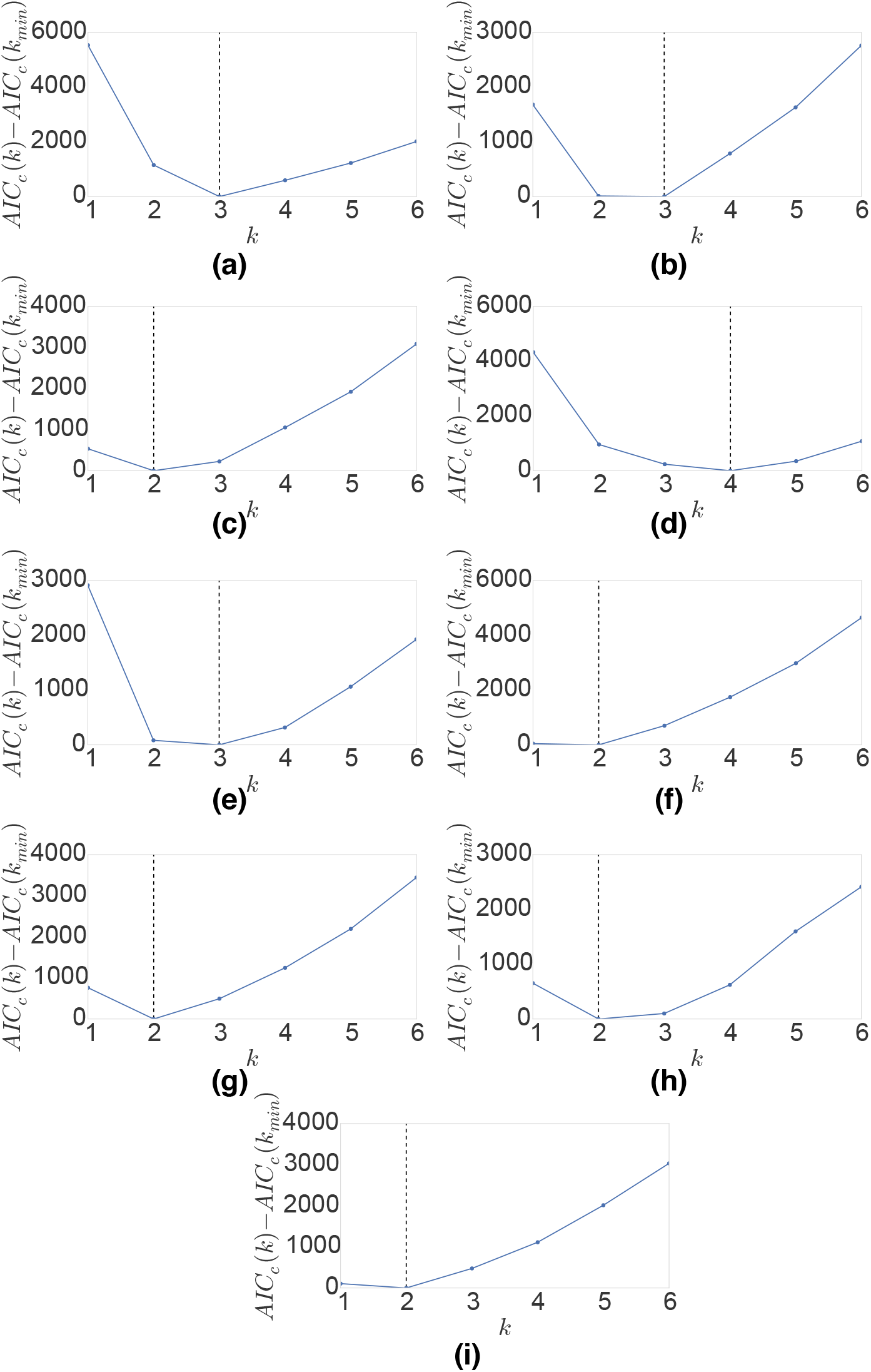
The best model is the one that minimizes Eq. 10, denoted as *AIC_c_*(*k_min_*), and consist of *K* syndromes. For each country we depict the relative likelihood of each candidate model (*AIC_c_*(*k*) − *AIC_c_*(*k_min_*)), where the *AIC_c_*(*k*) scores for each candidate model are compared against the AIC score of the best model *AIC_c_*(*k_min_*). We depict only models with k up to 6 and not 19 for easier visual inspection. The best model per country, with optimal number of syndromes is: (a) The Netherlands *K* = 3, (b) Belgium *K* = 3, (c) Italy *K* = 2, (d) France *K* = 4, (e) UK *K* = 3, (f) Spain *K* = 2, (g) Portugal *K* = 2, (h) Denmark *K* = 2, (i) Ireland *K* = 2. The best model is presented with dashed line.

**Supplemental Information, Figure S2.**
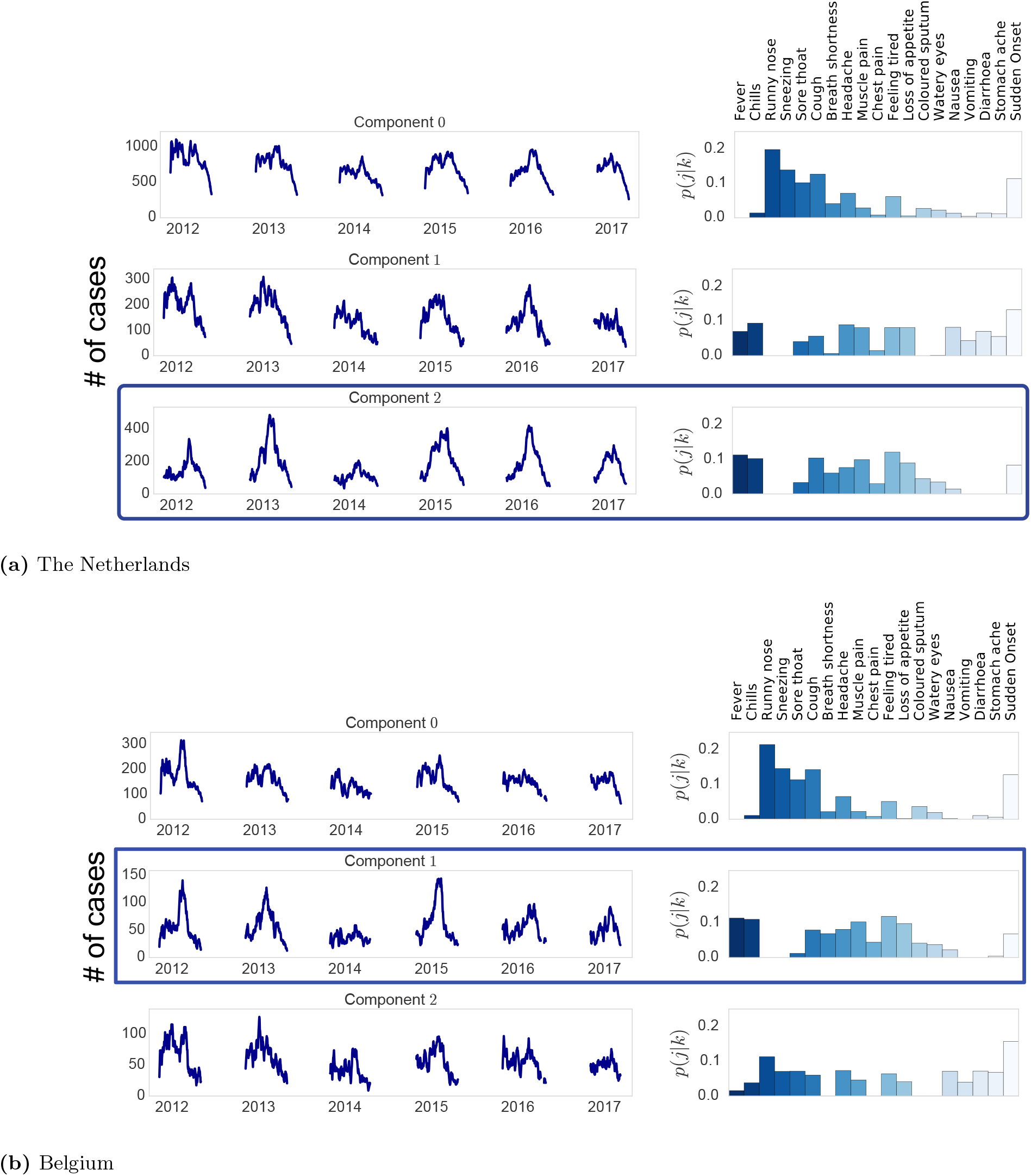
Comparative Analysis of the consistency and time series of the amount *y_ik_* which refers to the total number of counts associated to a syndrome *k* in day *i* for all the emerged syndromes for the Netherlands and Belgium. The blue box indicates the syndrome selected as IN_NMF by the algorithm.

**Supplemental Information, Figure S3.**
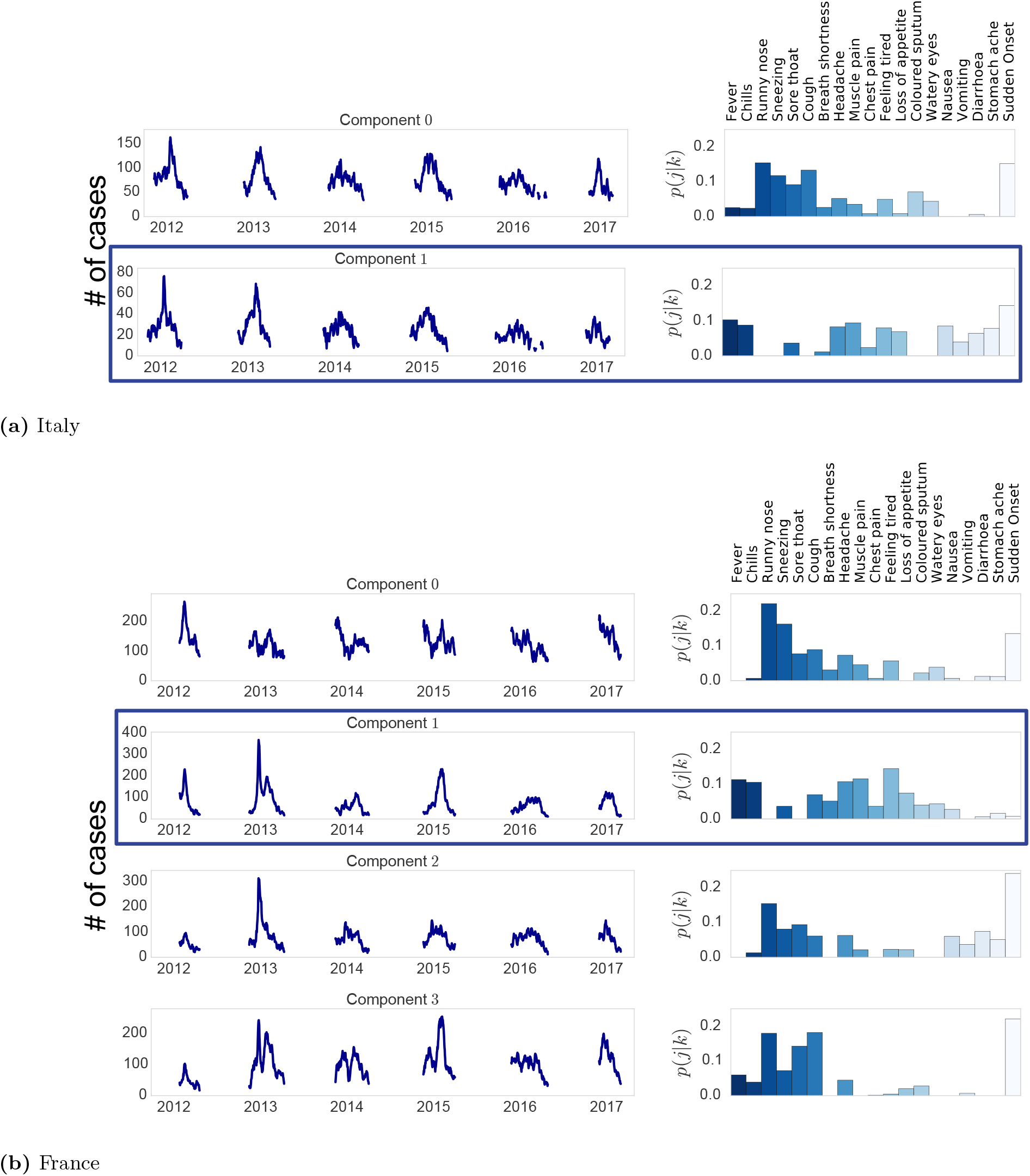
Comparative Analysis of the consistency and time series of the amount *y_ik_* which refers to the total number of counts associated to a syndrome *k* in day *i* for all the emerged syndromes for Italy and France. The blue box indicates the syndrome selected as IN_NMF by the algorithm. Note that for France the syndrome selected as IN _Gastro is the second component.

**Supplemental Information, Figure S4.**
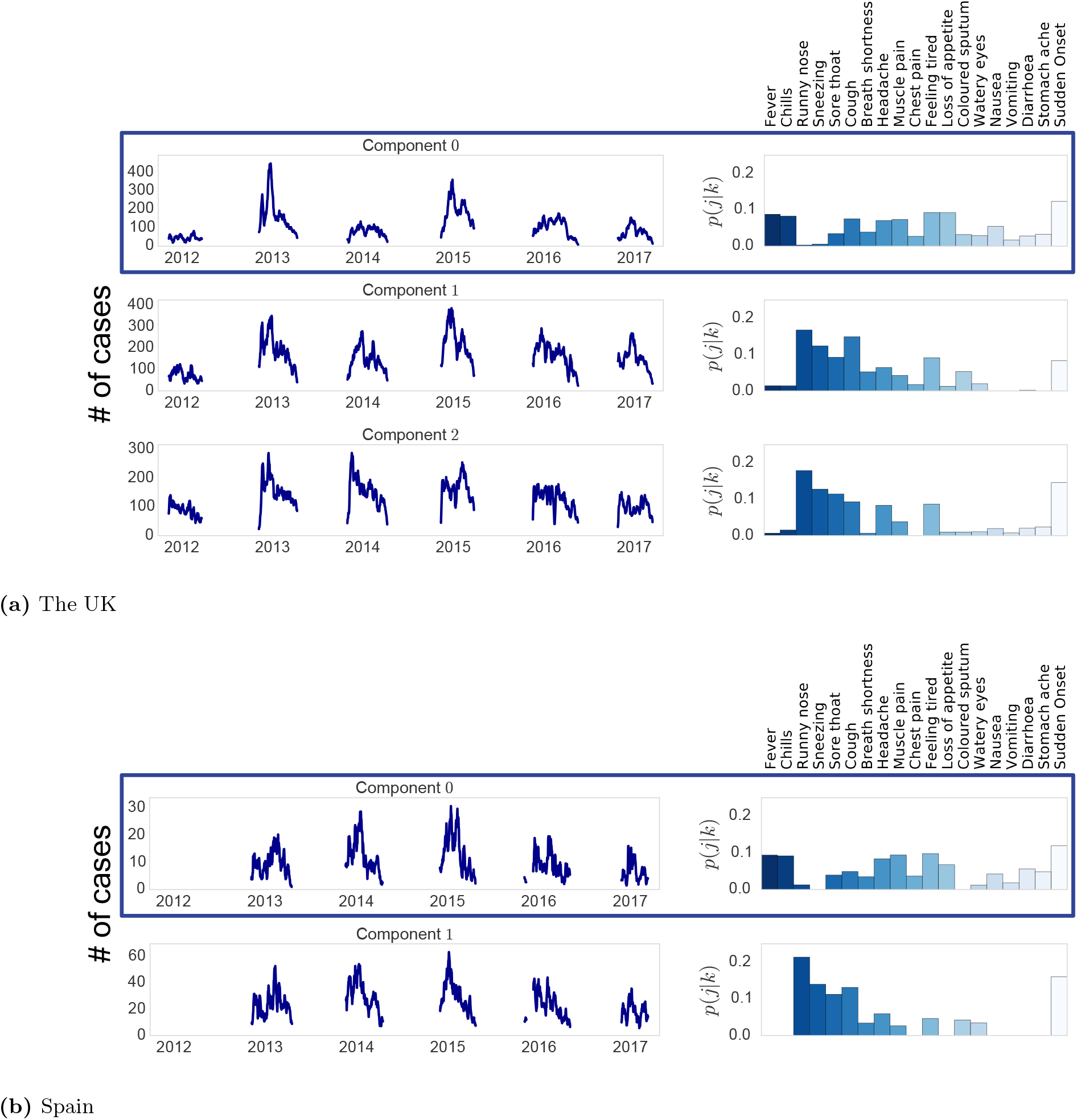
Comparative Analysis of the consistency and time series of the amount *y_ik_* which refers to the total number of counts associated to a syndrome *k* in day *i* for all the emerged syndromes for UK and Spain. The blue box indicates the syndrome selected as IN_NMF by the algorithm.

**Supplemental Information, Figure S5.**
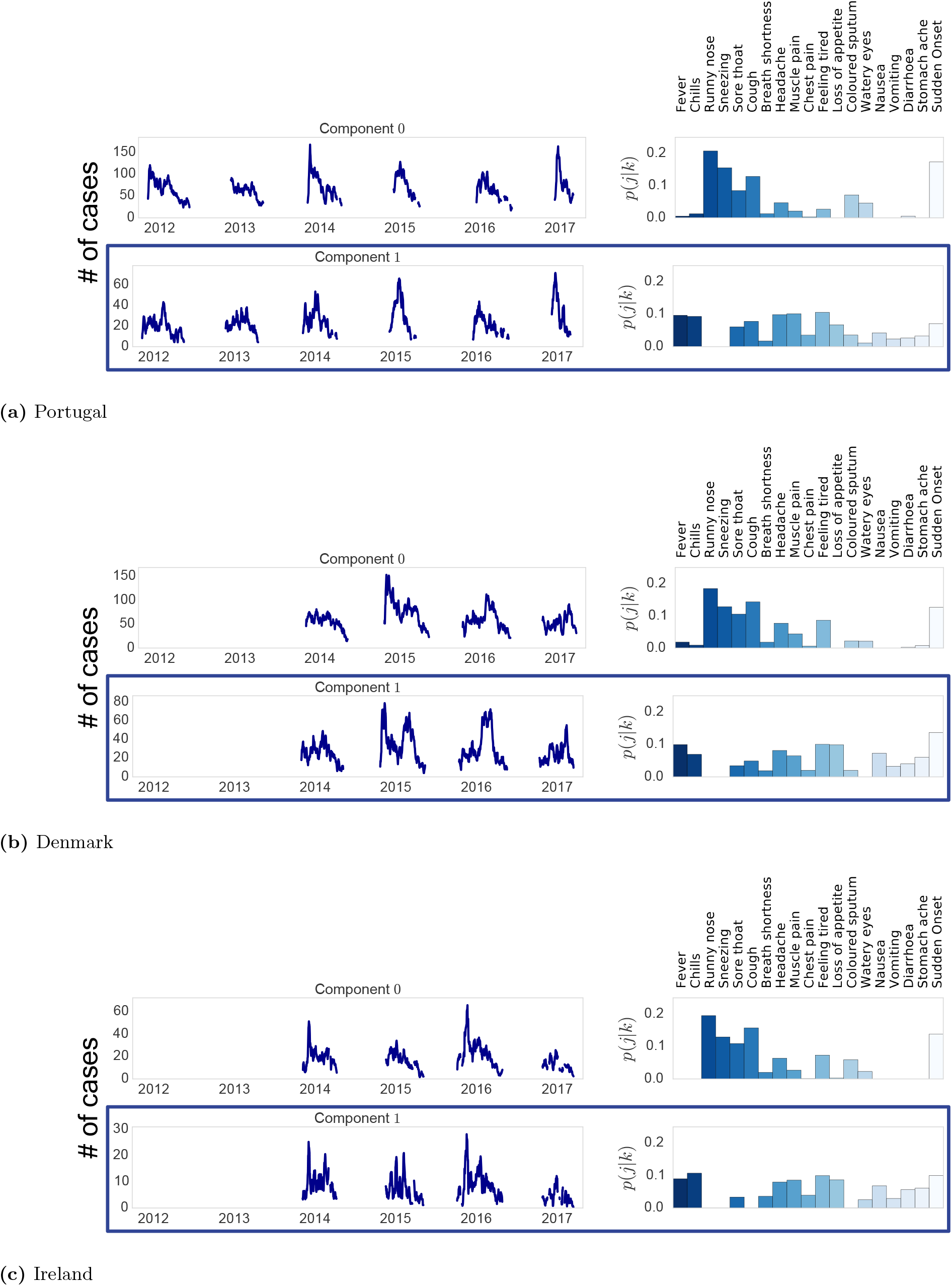
Comparative Analysis of the consistency and time series of the amount *y_ik_* which refers to the total number of counts associated to a syndrome *k* in the day *i* for all the emerged syndromes for Portugal, Denmark and Ireland. The blue box indicates the syndrome selected as IN_NMF by the algorithm. Note that for Denmark and Ireland we have data only for the period 2014 - 2017

## Footnotes

1 Throughout this study we employ the term *latent* as used in computer science, i.e. referring to variables that are hidden, not directly observed, but rather inferred through a mathematical model. There is no reference to the medical use of the term that usually indicates an asymptomatic infection. combination of symptoms as the best approximation for the actual influenza-like illness circulating among the general population.

2 We focused on the case of France due to the immediate data availability from the official surveillance. The *Réseau Sentinelles* in fact comprises a unique program of data collection about gastrointestinal illness episodes [48]

